# Diverse mechanisms for epigenetic imprinting in mammals

**DOI:** 10.1101/2021.04.30.442087

**Authors:** Daniel Andergassen, Zachary D. Smith, John L. Rinn, Alexander Meissner

## Abstract

Genomic imprinting and X chromosome inactivation (XCI) require epigenetic mechanisms to direct allele-specific expression. Despite their critical roles in embryonic development, how universal epigenetic regulators coordinate these specific tasks from single locus to chromosome-scale remains understudied. Here, we systematically disrupted multiple essential epigenetic pathways within polymorphic F1 zygotes to examine postimplantation effects on canonical and non-canonical genomic imprinting as well as X chromosome inactivation. We find that DNA methylation and Polycomb group repressors are both indispensable for autosomal imprinting, albeit at distinct gene sets. Moreover, the extraembryonic ectoderm relies on a broader spectrum of unique imprinting mechanisms, including non-canonical targeting of maternal endogenous retrovirus (ERV) driven promoters by G9a. We further utilize our data to identify Polycomb dependent and independent gene clusters on the imprinted X chromosome, which appears to reflect distinct domains of Xist-mediated suppression. Our data has allowed us to assemble a comprehensive inventory of the epigenetic mechanisms utilized in eutherian mammals to maintain parent-specific imprinting, including an expanded view of the placental lineage that comprises multiple unique pathways.

## INTRODUCTION

Mammals have two autosomal gene copies, one inherited from each parent. The vast majority of genes are biallelically expressed, while a small subset is expressed in a parent of origin-specific fashion. Genomic imprinting is maintained via distinct epigenetic mechanisms to propagate information from the oocyte or sperm into the next generation, and can regulate the expression of nearby genes in *cis* as a secondary mechanism. Multiple loci have been thoroughly dissected to understand how this process is carried out molecularly. For example, epigenetic modifications can repress long non-coding RNAs (lncRNAs) that otherwise target and repress nearby genes. For decades, DNA methylation was considered the only epigenetic modification that could be transmitted from the germline into the subsequent progeny. Recently, oocyte-specific trimethylation of H3 on lysine 27 (H3K27me3) was found to transiently imprint several loci within preimplantation embryos, of which some transition to a more permanent DNA methylated state to preserve paternal expression (Chen et al., 2019; Hanna et al., 2019; Inoue et al., 2017a). Non-canonical imprinting by H3K27me3 appears to primarily function in placental development and regulates genes with critical functions in this lineage. This surprising alternative imprinting strategy implies that mammals may use other mechanisms beyond DNA methylation to instruct parent-specific regulation.

Notably, oocyte-specific H3K27me3 also serves as a maternal imprint for the lncRNA *Xist*, which triggers paternal X chromosome inactivation (XCI) in mouse preimplantation embryos and extraembryonic tissues (Inoue et al., 2017b). While similar to non-canonical H3K27me3-based regulation of single loci, X inactivation is capable of relaying this information to suppress an entire chromosome and also operates through several classic epigenetic suppression pathways (Żylicz and Heard, 2020). Failure to establish or maintain either XCI or genomic imprinting results in embryonic lethality, emphasizing the developmental importance of these interrelated processes. However, the molecular relationship between imprinted XCI and other H3K27me3 regulated loci, or what distinguishes these strategies from canonical, DNA methylation-based imprinting, remains unresolved.

To compare and contrast the developmental roles of epigenetic regulators in these processes, we used reciprocal crosses between highly polymorphic strains to investigate allele-specific expression in E6.5 epiblast (the embryonic progenitor) and extraembryonic ectoderm (ExE, the placental progenitor, which exhibits imprinted XCI). With these data, we generate a comprehensive map of all the imprinted genes in both embryonic and extraembryonic lineages, as well as genes that escape imprinting on the inactive X chromosome. Then, we systematically perturbed multiple epigenetic pathways, including DNA methylation, Polycomb-based repression, and Histone 3 Lysine 9 methylation (H3K9me), to understand their primary contributions to parent-specific expression. We find that DNA methylation appears to function primarily at previously described canonical Imprint Control Regions (ICRs) within both lineages, whereas the early placenta exhibits a greater diversity of imprinting mechanisms. For example, we identify an H3K9 methylation-based mechanism that recognizes and suppresses ERV-driven promoters exclusively at maternal loci. Furthermore, our observations pinpoint the explicit dependencies of distinct X chromosomal regions on the Polycomb repressor complexes (PRCs), including gene clusters that appear to rely on *Xist* recruitment alone. We also find that PRC2 has independent roles in *Xist* imprinting and chromosome-wide silencing. Finally, our data set enables us to generate a complete inventory of parent-specific expression signatures as they depend on epigenetic pathways.

## RESULTS

### Capturing allele-specific expression in the early embryo and placenta

To explore parent-specific expression within the early embryonic and extraembryonic lineages, we conducted reciprocal crosses between CAST/EiJ (CAST) and B6D2F1/J (BDF1) strain animals, isolated embryonic day (E)6.5-stage epiblast (Epiblast) and extraembryonic ectoderm (ExE), and performed RNA-seq (**Figure 1A, Table S1 sheet A**). On average, this approach allowed us to call allele-specific expression for 15,199 and 13,729 genes for the Epiblast and ExE, respectively, varying between 14,136–20,567 and 13,500-14,143 genes per replicate (informative gene ≥ 10 reads SNPs overlap, **see Methods**). Unsupervised clustering of gene expression confirmed epiblast and ExE lineage identity and sample purity. Overall, allelic ratios exhibited expected patterns: biallelic expression for autosomal genes, skewed XCI that favors expression of the CAST allele in Epiblast (Calaway et al., 2013), and imprinted XCI in ExE (**Figure S1A-B**). For autosomal imprinting, we identified 22 Epiblast- and 41 ExE-specific imprinted genes, with 19 shared between lineages (**Figure 1B-D, Table S1 sheet B-C**). We also detected the seven known non-canonical imprinting genes (*Gab1, Sfmbt2, Slc38a4, Phf17, Smoc1, Platr20*, and *Gm32885*) in ExE, as well as seven putative, novel imprinted genes (Epiblast *n* = 1 and ExE *n* = 6). Our newly discovered imprinted loci are located in proximity to known imprinted regions and include six imprinted lncRNAs as well as maternally dominant expression of Brachyury (*T*) and *Pnldc1*.

**Figure 1.**
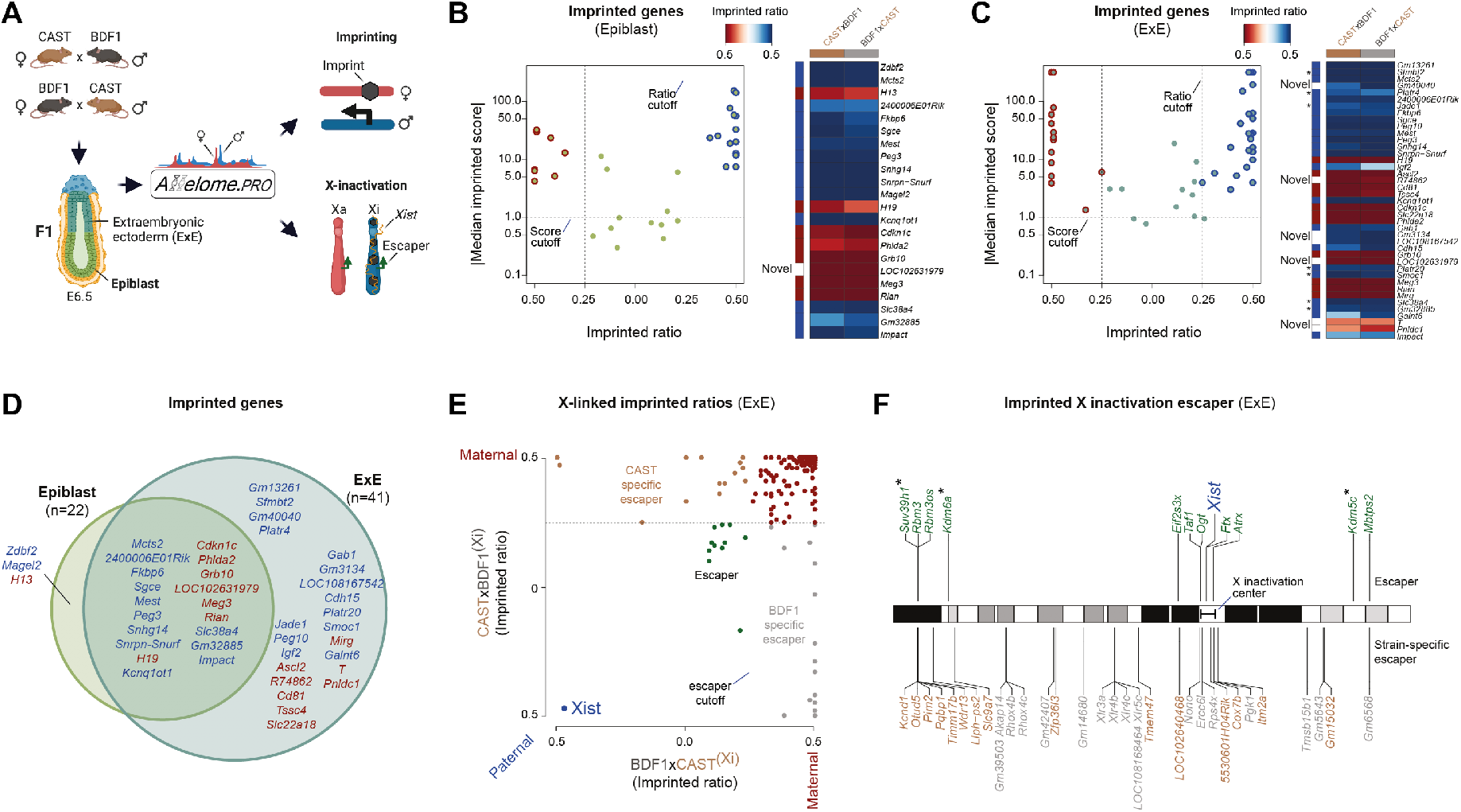
An inventory of parental-specific expression for male and female embryos in early embryonic and extraembryonic lineages. (A) Simplified schematic of the experimental system used to obtain a parent-specific expression landscape (imprinting and X-inactivation). E6.5 Epiblast (light green) and extraembryonic ectoderm (ExE, dark green) are isolated from F1 reciprocal crosses (*n* = 11, BDF1xCAST 4m/3f and CASTxBDF1 1m/3f) and subjected to RNA-seq. Colors are used in figures throughout the manuscript to highlight the tissue of origin. (B-C) Imprinted genes identified in the E6.5 Epiblast and ExE (red: maternally expressed, blue: paternal expressed) using a median imprinted score cutoff and allelic ratios of 1 and 0.25, respectively (dashed line). Corresponding heatmaps show the imprinted ratios in the forward (BDF1xCAST) and reverse (CASTxBDF1) cross. Novel identified imprinted genes are indicated in white. This also includes the observation of maternal-specific expression of the major regulator Brachyury (*T)* within the ExE. (D) Overlap of imprinted genes between E6.5 Epiblast and ExE. (E) Scatter plot showing the imprinted ratios for X-linked genes in ExE between forward (BDF1xCAST) and reverse (CASTxBDF1) crosses. Maternally expressed genes (red), XCI escaper genes (green), and strain-specific escape from the CAST (brown) and BDF1 (grey) inactive X chromosome are indicated. The dashed line indicates the escaper cutoff of 0.25. (F) Chromosomal overview of genes that maintain biallelic expression on the Xi (“escapers”) with shared above and strain-specific below. Asterisk indicates escaper genes that function as chromatin modifiers.

To explore chromosome-level regulation, we next examined expression of X-linked genes in female ExE samples, which undergo imprinted XCI (**Figure 1E, Table S1 sheet D**). Of 335 informative X-linked genes, 290 (86.5%) are maternally expressed, 11 (3.3%) escape inactivation (‘escaper,’ excluding *Xist*), and 33 (9.85%) escape X inactivation in a strain-specific manner (Cast *n* = 15, and BDF1 *n* = 18). Half of escaper genes cluster in proximity to the 1.85-Mb X inactivation center candidate region defined in Chadwick *et al*. (Ref (Chadwick et al., 2006)), while others include notable epigenetic regulators (e.g., *Suv38h1, Kdm6a, Kdm5*c, **Figure 1F**). In summary, we mapped the parental-specific transcriptional landscape in the early embryonic and extraembryonic lineage for both males and females, allowing us to examine autosomal and X-chromosome-specific imprinting.

### Determining imprinting regulators by zygotic genome perturbation

Our comprehensive map of parent-specific expression allows us to systematically investigate the roles of key epigenetic pathways by Cas9-based genetic disruption in zygotes (Grosswendt et al., 2020; Smith et al., 2017) (**Figure 2A, see Methods**). For this study, we disrupted the DNA methyltransferase Dnmt1, the histone 3 lysine 9 (H3K9) methyltransferases G9a and GLP (target genes *Ehmt1* and *Ehmt2*, double KO), as well as the PRC1 and PRC2 complexes individually (target genes: *Rnf2* and *Eed*, respectively) in BDF1xCAST zygotes. Epiblast and ExE samples were collected from 30 knockout embryos at E6.5 and processed for bulk RNA sequencing (**Table S1 sheet E-F**). We validated gene disruption by comparing the expression profile and level of the target gene between wildtype and KO embryos (**Figure S2A-B**) and excluded imprinted genes with less than two informative allelic ratios in any regulator disruption, resulting in 19 epiblasts and 37 ExE imprinted genes. Next, we performed unsupervised clustering of autosomal expression levels and confirmed all samples preserved their overall lineage identity (**Figure S2C**). Within the Epiblast and ExE clusters, we found that *ΔDnmt1* and *ΔG9a-GLP* embryos co-cluster, as do *ΔRnf2* and *ΔEed*, in keeping with their more similar functions (Auclair et al., 2016; Jiang et al., 2020).

**Figure 2.**
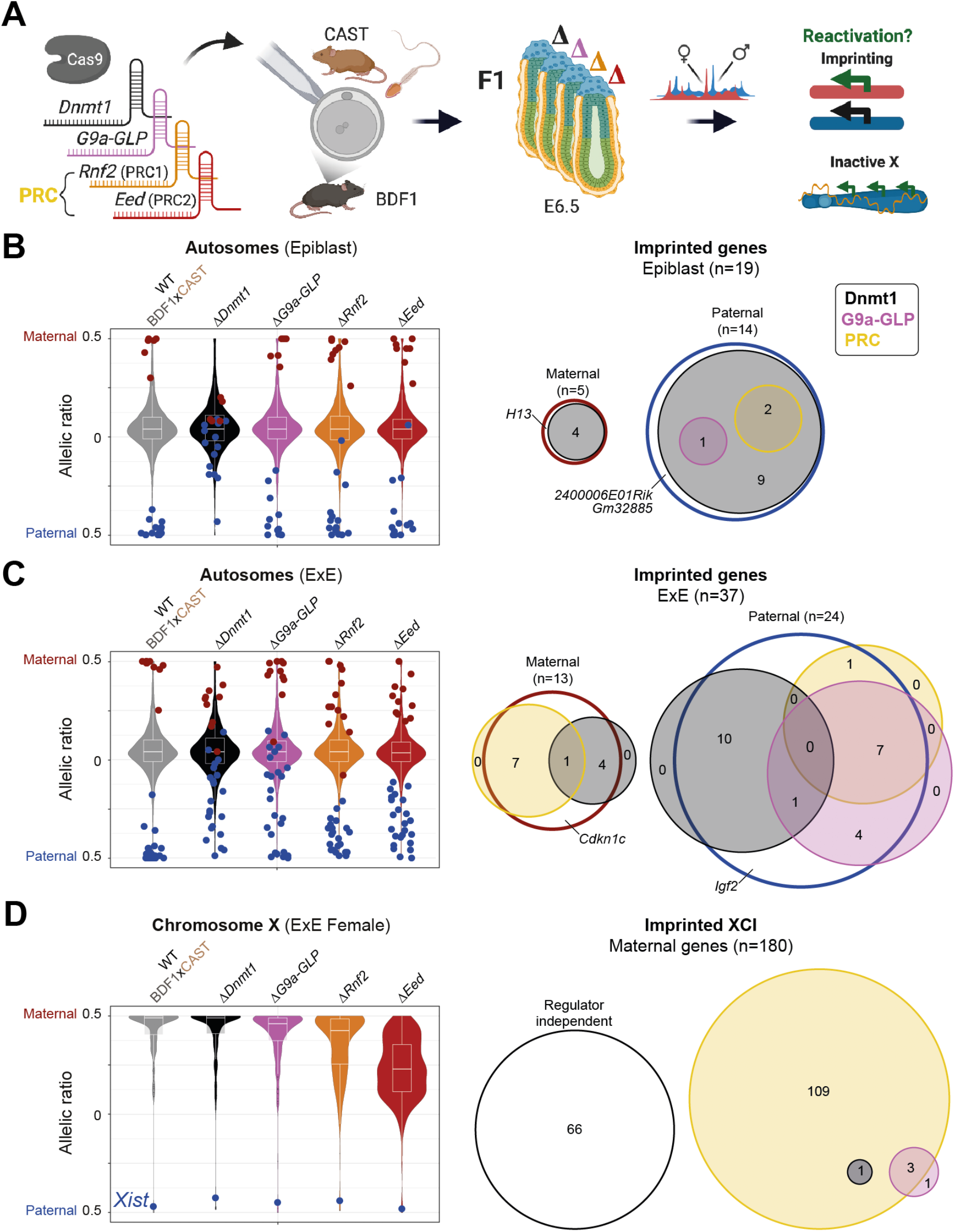
Epigenetic regulation of autosomal and X chromosome-specific imprinting. (A) Schematic overview for our strategy to assign roles for selected epigenetic key regulators to parent-specific gene expression in E6.5 Epiblast (light green) and extraembryonic ectoderm (ExE, dark green). Specific epigenetic regulators are disrupted by injection of Cas9 and sgRNAs into hybrid F1 (BDF1xCAST) Zygotes (*ΔDnmt1 n =* 3, *ΔG9a-GLP n* = 9, *ΔRnf2 n* = 8, *ΔEed n* = 10). Regulator colors are used throughout the rest of the manuscript: *Dnmt1*, black; *G9a* and *GLP*, violet, gene name *Ehmt1* and *Ehmt2*; PRC1 member *Rnf2*, orange; and PRC2 member *Eed*, red. (B) Violin plots of the median allelic-ratios of autosomal genes from WT and regulator knockouts in the epiblast (left). Maternal and paternal expressed imprinted genes are indicated with red and blue dots, respectively. Venn diagram shows the intersection of each epigenetic regulators’ contribution in maintaining imprinted genes in the epiblast (right). An epigenetic regulator was counted as relevant for silencing imprinted genes if the change in allelic ratio between WT and KO was ≥ 20%. PRC1 and PRC2 were summarized as PRC by using the higher delta. DNA methylation-dependent imprinting is most frequent in the epiblast. Regulator independent imprinted genes have a delta allelic ratio < 20% in all disrupted regulators. Imprinted genes with less than two informative allelic ratio values in any regulator disruption dataset are not shown. (C) As in (B) for the ExE lineage. Extraembryonic imprints appear to depend on a more diverse set of regulators. (D) Violin plots displaying the median allelic-ratio of X-linked genes from WT and regulator knockouts in female ExE (left). The blue dot highlights the allelic ratio of the lncRNA *Xist*. Successful extraembryonic XCI depends on PRC2 and, to a lesser degree, PRC1 downstream of paternal *Xist* expression, which is stable in all regulator mutants examined (**Figure S2D**). Venn diagram for imprinted XCI (right) as shown for autosomal imprinting in (B-C). An epigenetic regulator was counted as relevant for silencing X-linked genes if the delta allelic ratio change between WT and KO for any regulator was ≥ 20%. Regulator independent X-linked genes have a delta allelic ratio < 20% in all disrupted regulators.

### The extraembryonic lineage uses an expanded set of imprinting mechanisms

To test which epigenetic regulators are responsible for parent-specific regulation, we first examined the allelic landscape of autosomes. We observed biallelic expression of imprinted silencing for most identified alleles within the *ΔDnmt1* Epiblast, the majority of which are also deregulated in ExE (**Figure 2B, Table S1 sheet G**). In contrast, the ExE includes several additional dependencies that largely reflect parent of origin and epigenetic mechanism. For example, G9a appears to regulate a discrete set of paternally imprinted genes, which also depend to a lesser degree on PRC regulators. In contrast, *Rnf2* and *Eed* mutants preferentially influence the expression of maternal imprints and do so independently of either H3K9 or DNA methylation (**Figure 2C, Table S1 sheet H**). After examining our mutant data, only five imprinted genes (three in Epiblast and two in ExE) cannot be explained by at least one of these key pathways and remain imprinted across our mutant cohorts (delta allelic ratio < 20%).

We also find that the PRCs stabilize imprinted X inactivation, with PRC2 disruption more severe than PRC1 (**Figure 2D, Figure S2D**). Loss of PRC2 regulation during XCI mirrors the response of PRC2-dependent imprints, with muted reactivation of the silenced allele or paternal chromosome, respectively. Notably, we do not observe disrupted *Xist* imprinting or changes in allelic expression for ∼36% of X-linked genes across all mutant regulators examined. Together, the preserved imprinting of *Xist* and the muted reactivation of paternal alleles suggest that PRC2 acts to stabilize subsets of *Xist* lncRNA target genes, an observation we explore in greater detail below.

### G9a-GLP control a discrete set of non-canonical imprinted genes

In Epiblast, the majority of imprinted genes appear to depend primarily on DNA methylation (*n* = 14 out of 19 genes, 3 of which do not change in any KO, **Figure 3A left, Figure S3A**). *Zdbf2* and *Slc38a4* are notable exceptions: *Zdbf2* is reactivated in both *Rnf2* and *Eed* mutants, whereas *Slc38a4* primarily relies on G9a-GLP, in line with previous reports (Auclair et al., 2016; Greenberg et al., 2017). Notably, most epiblast-associated imprinted regions exhibit similar changes and dependencies in the ExE, suggesting that these are constitutive and do not depend on their respective lineage (**Figure 3A right, Figure 3B, Table S1 sheet I**). In contrast, 20 out of 37 total imprinted genes in the ExE are DNA methylation independent (**Figure 3A right**). Of these, seven maternally expressed loci are chromatin-dependent and predominantly rely on PRC1 and 2 (**Figure 3B top**). All of these are either known targets or in local proximity to paternally expressed lncRNAs *Kcnq1ot1* (*Slc22a18, Alscl2, Tssc4, R74862, Cd81*) and *Airn (T, Phlda2)*, which are believed to recruit these complexes to enable silencing of paternal loci (Andergassen et al., 2017, 2019; Pandey et al., 2008; Schertzer et al., 2019; Terranova et al., 2008; Vausort et al., 2014).

**Figure 3.**
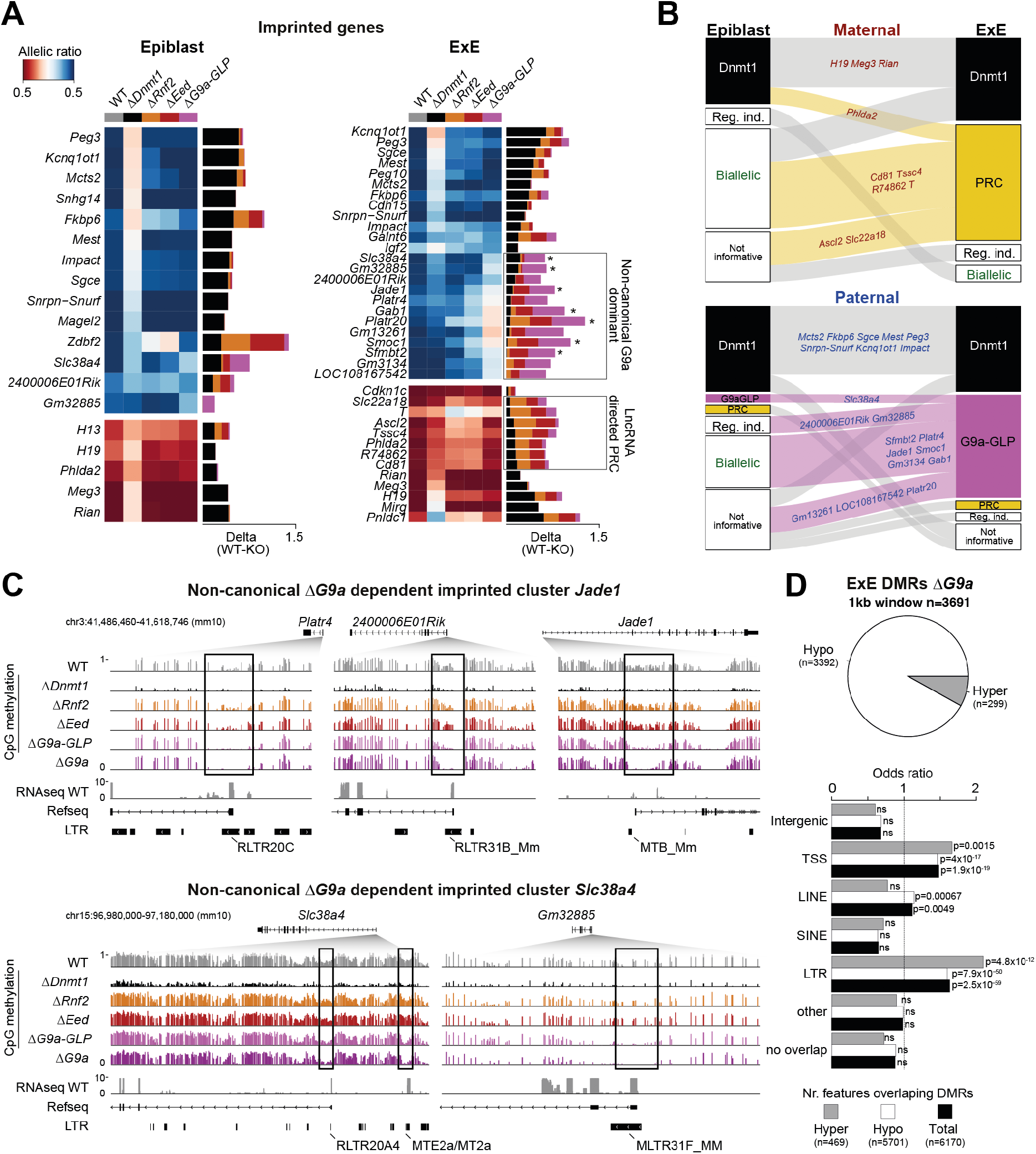
G9a controls endogenous retrovirus promoters of non-canonical imprinted genes. (A) Allelic ratio of imprinted genes and the corresponding delta changes between wildtype (BDF1xCAST) and regulator disruption identified in the Epiblast (left) and ExE (right) lineage. Heatmap ranked by delta allelic ratio change between WT and *Dnmt1* deletion. For ExE, boxes highlight DNA methylation independent non-canonical expressed genes (known* and novel) and lncRNA directed PRC targets. Notably, ExE-specific imprinting is most apparent for a set of paternally-expressed, G9a-GLP controlled loci that are only weakly dependent on PRCs. Imprinted genes with less than two informative allelic ratio values in any regulator disruption dataset are not shown. (B) Flow diagram outlining changes in the imprinted landscape between Epiblast and ExE for maternally (top) and paternally (bottom) expressed genes. (C) Genome browser tracks for WGBS and RNA-seq data from WT and regulator disrupted E6.5 ExE lineages capturing the two non-canonical G9a dependent imprinted clusters *Jade1* (top) and *Slc38a4* (bottom). Boxes highlight G9a dependent hypomethylated DMRs (Overlapping ERV LTRs are indicated). (D) Identified ExE DMRs using WT and *ΔG9a* WGBS data (1kb window, *n* = 3691, |delta cutoff| ≥ 20%). Pie chart showing the proportion of hypo- and hypermethylated DMRs (top). Feature enrichment of the identified DMRs over background was calculated for intergenic, genic (±1kb of TSS), and different repeat classes using the Fisher’s exact test (bottom).

Notably, half of paternally expressed, ExE-specific imprinted genes appear to depend primarily on G9a-GLP (G9a-GLP *n =* 12, Dnmt1 *n* = 11, PRC2 *n* = 1), which include all previously defined H3K27me3-dependent non-canonical imprinted genes (Chen et al., 2019; Hanna et al., 2019; Inoue et al., 2017a) (**Figure 3A right, Figure 3B bottom, Table S1 sheet J**). These results show an interaction between H3K9 and H3K27 methylation and suggest that H3K9me may be the dominant modification for maintaining and possibly establishing non-canonical imprinting. We also see some indication that these non-canonical imprints are distributed in clusters. For example, genes such as *Platr4 and 2400006E01Rik* are close to the known non-canonical imprinted gene *Jade1*, suggesting that this pathway may control an entire imprinted cluster on chromosome 3 (**Figure 3C top**). G9a-GLP dependent cluster regulation was also observed for the non-canonical imprinted genes *Slc38a4* and *Gm32885* on chromosome 15 (**Figure 3C bottom**). Finally, G9a-GLP regulation was also recognized for the paternally expressed *Sfmbt2* gene, including one of the genome’s largest miRNA clusters.

### G9a recognizes ERV-driven promoter elements

A recent study reported that the long terminal repeats (LTR) of ERV elements can act as alternative promoters for non-canonical imprinted genes and found that maternally-inherited H3K27me3 transitions to DNA methylation-based silencing in extraembryonic tissues (Hanna et al., 2019). To examine the effects of G9a-GLP on these elements, we examined the DNA methylation dynamics of our G9a-sensitive genes using published whole-genome bisulfite sequencing (WGBS) data in regulator disrupted E6.5 ExE *(ΔDnmt1, ΔRnf2, ΔEed* and *ΔG9a*) (Grosswendt et al., 2020) as well as newly generated data for our *G9a-GLP* double KO (**Figure 3C**). Strikingly, all of our identified non-canonical imprinted genes show local loss of DNA methylation around their transcription start sites (TSSs) in *G9a* KOs, but not for either PRC mutant (**Figure 3C, Figure S3B**). Furthermore, within these imprinted loci, every G9a-dependent differentially methylated region (DMR) overlaps an ERV LTR (**Figure 3C, Figure S3B**). This finding further strengthens our prior RNA-seq based observation that H3K9me appears crucial for non-canonical imprinting and suggests that G9a alone is sufficient to recruit DNA methylation in this context. Next, we investigated whether G9a regulation of ERV-driven promoters is a general regulatory mechanism or specific for non-canonical imprinted regions. To address this question, we first defined ExE DMRs using WT and ΔG9a WGBS data (**see Methods**) and identified 3392 hypomethylated (91.8%) and 299 hypermethylated (8.1%) DMRs (**Figure 3D, Table S1 sheet K**). The highest enrichment for hypomethylated DMRs over background was observed for promoters (*P* = 4×10^−17^, odds-ratio=1.5, +/- 1kb TSS) and LTRs (*P* = 7.9×10^−50^, odds-ratio=1.6). In summary, we find that G9a is the critical regulator for ERV-driven non-canonical imprinting and propose that this could be a more general gene regulatory mechanism.

### Epigenetic regulation of the inactive X chromosome in females

We next leveraged our data to explore the architecture of X inactivation as it is regulated by distinct epigenetic pathways. Out of 180 maternally expressed X-linked ExE genes (informative in all investigated ExE samples), 114 (63.4%) change their allelic expression in at least one epigenetic regulator mutant. These are almost entirely explained by PRC-based regulation, for which PRC2 is dominant: 74 X-linked genes are derepressed exclusively in Δ*Eed* embryos (65%), while another 35 genes are shared between Δ*Rnf2* and Δ*Eed* (30.7%) (**Figure 4A**). This synergy between PRC1 and PRC2, together with previous reports that maternal EED is sufficient to initiate and establish imprinted XCI (Harris et al., 2019), confirms that Polycomb-based repression is critical for propagating chromosome-wide epigenetic suppression. To learn more about the remaining 66 regulator independent genes (36.6%), we plotted allelic ratios according to their genomic position across the X chromosome. PRC-independent genes exist in defined clusters and appear to be independent of WT expression levels (**Figure 4B, Figure S4A**). We, therefore, hypothesized that these clusters might represent initial *Xist* target loci, for which imprinted *Xist* expression (and localization) is sufficient to maintain paternal silencing. To explore this, we investigated published *Xist* RNA Antisense Purification (RAP) data, a method that maps lncRNA interactions with chromatin (Engreitz et al., 2013) as well as H3K27me3 and H2AK119ub ChIP-seq specific for the inactive X chromosome (Żylicz et al., 2019). Our analysis shows a strong enrichment for *Xist* binding, H3K27me3, and H2AK119ub at PRC-independent loci and an inverse correlation between direct *Xist* binding and PRC dependence (**Figure 4C**). Taken together, these findings highlight a central role for both PRCs in translating local *Xist* recruitment to stably silence the surrounding area.

**Figure 4.**
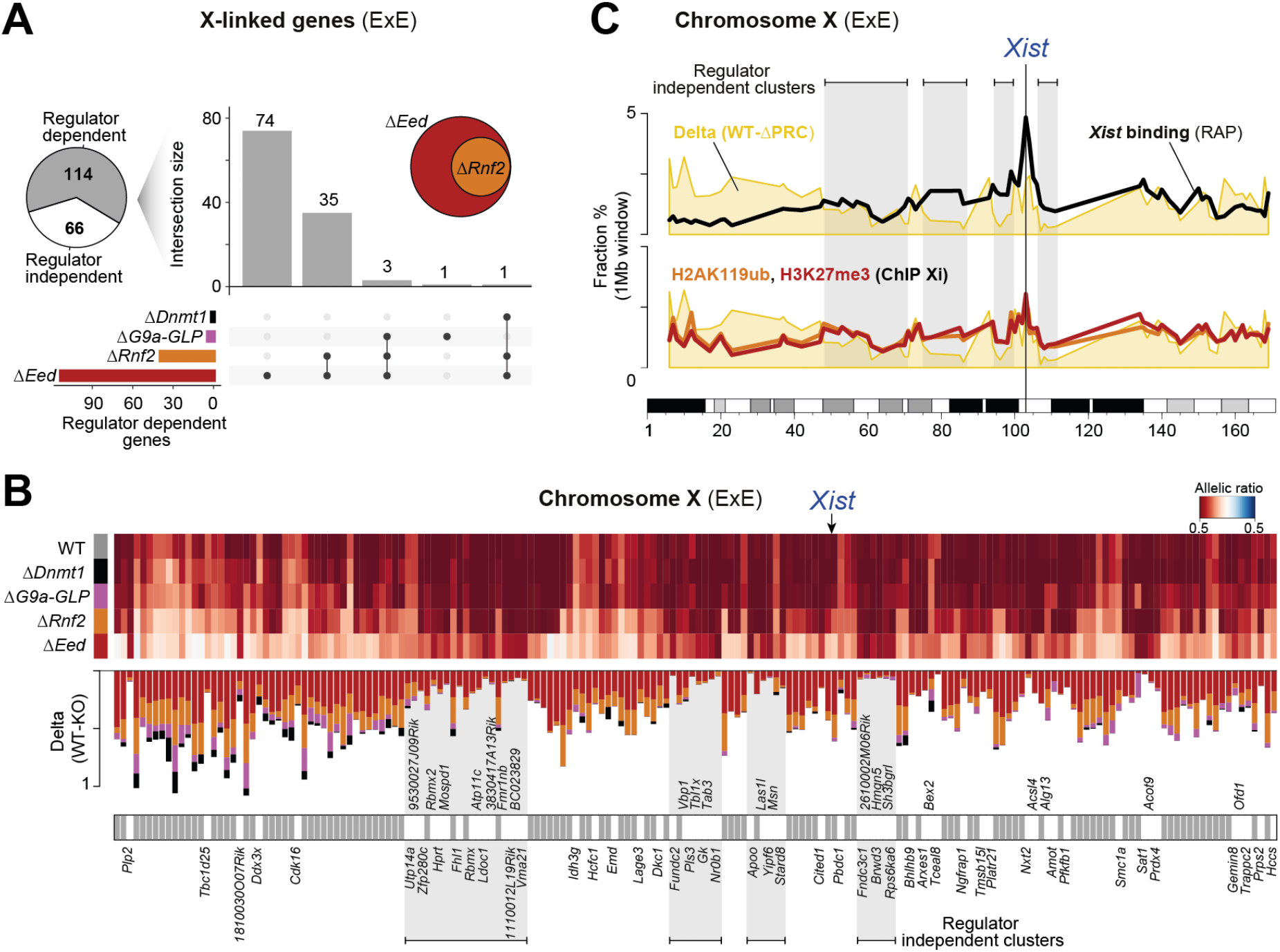
Polycomb-based repression is critical for the maintenance of the imprinted X chromosome. (A) Pie chart showing the proportion of regulator dependent or independent genes for imprinted XCI. A gene was called regulator dependent if the delta allelic ratio changes between WT and KO ≥ 20% for any regulator. UpSet plot shows the overlap across the four disrupted regulator members: *Dnmt1, G9a-GLP, Rnf2*, and *Eed*. The inset Venn diagram highlights the high overlap between *Eed* and *Rnf2* dependent X-linked genes. Genes that escape imprinted XCI were excluded from this analysis (B) Regulator-independent X-linked genes are organized into distinct spatial clusters. Allelic ratios of maternally expressed X-linked genes and the corresponding delta changes between wildtype (BDF1xCAST) and regulator disrupted E6.5 ExE lineages are shown (X-linked gene Matrix ranked by the genomic position). Regulator-independent X-linked genes are indicated. A gene was called regulator independent if the delta allelic ratio changes between WT and KO < 20% for every regulator. (C) Regulator independent regions occur in domains with high *Xist* enrichment and repressive chromatin. 1Mb windows summarizing (average) the X-linked gene delta changes between WT and PRC, for Xist enrichment over input (RAP) (Engreitz et al., 2013) and for H3K27me3 and H2AK119ub enrichment on the inactive X (Żylicz et al., 2019) across the entire X-chromosome.

### Three distinct mechanisms for encoding parental-specific gene regulation

Our results allowed us to build a complete inventory of genes that exhibit parental-specific expression in the early embryo and placenta and evaluate their dependencies on key epigenetic pathways. From our perturbation data, we are able to assign the regulation of every imprinted region in Epiblast (*n* = 13) and ExE (*n* = 20) to one of three distinct epigenetic mechanisms (**Figure 5A**). First, known canonical ICRs are established and maintained by germline DMRs. Second, paternally unmethylated ICRs enable lncRNA transcription, which then recruits repressive machinery to distal target genes in *cis*. Third, a G9a dominant mechanism controls ERV-driven non-canonical imprinting in extraembryonic tissues at genes that are often clustered together. Of the thirteen regulator-dependent imprinted regions in the embryonic lineage, all are associated with gDMRs and primarily represent ICRs (**Figure 5B**): eleven regions are directly linked to gDMRs, while two regions, *Zdbf2* and *Slc38a4*, translate gDMR information *in cis* via PRC2, with G9a acting as a secondary mechanism (Auclair et al., 2016; Greenberg et al., 2017). Lastly, the *Kcnq1* region directs DNMTs via the lncRNA *Kcnq1ot1* to silence the distant gene *Phlda2*, in line with a previous report (Mohammad et al., 2010).

**Figure 5.**
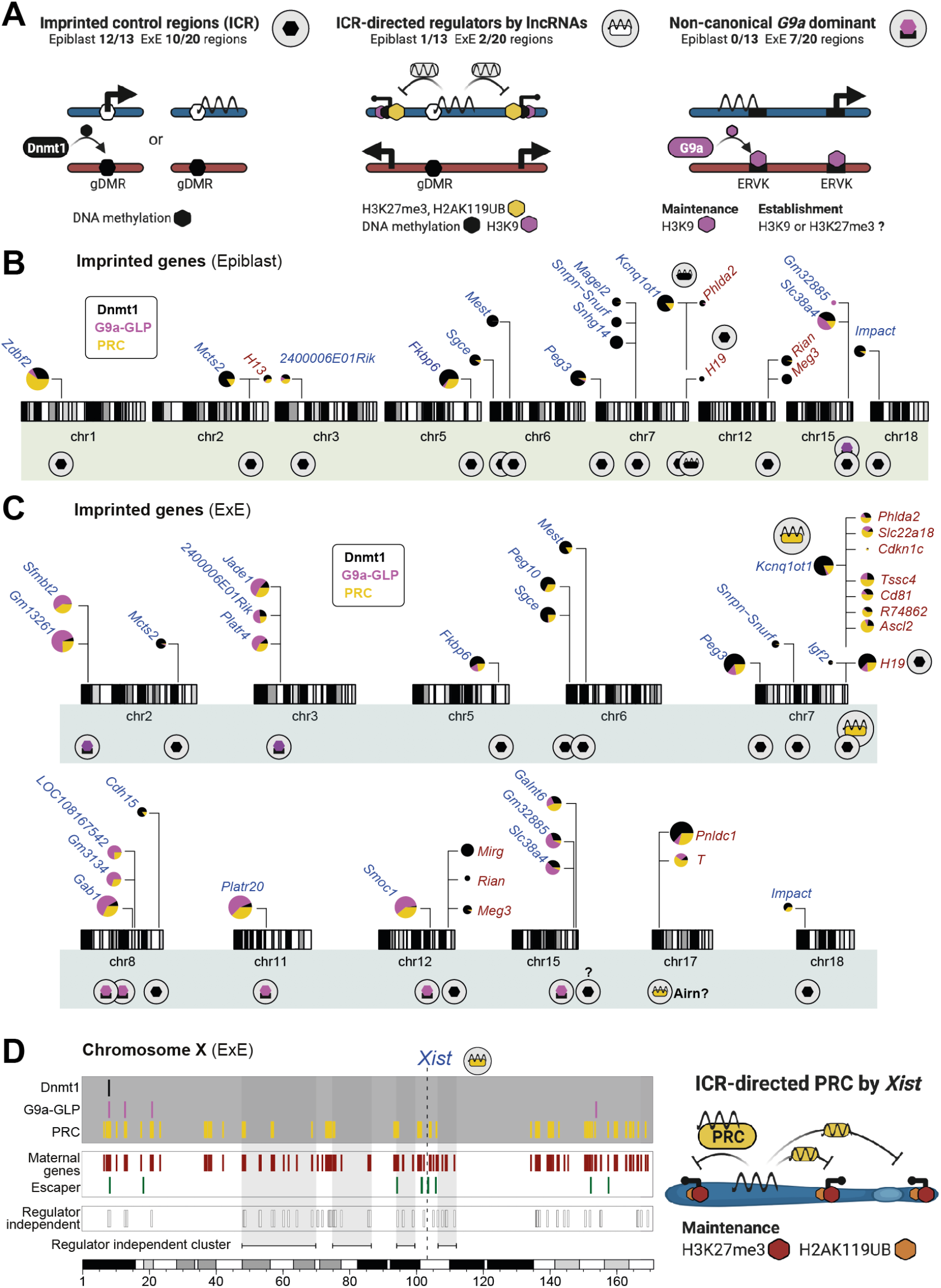
Three diverse epigenetic mechanisms for controlling parent-specific gene regulation. (A) Illustration of the identified mechanisms for parental-specific gene regulation: Imprinted control region (ICR), ICR-directed lncRNA deployment, and non-canonical G9a dominant. (B) Ideograms of mouse chromosomes show the position of Epiblast-specific imprinted genes. Pie charts on the top highlight the proportion of the delta allelic ratio change between WT and the disrupted regulators *Dnmt1, G9a-GLP*, and PRC. The circle size denotes the combined delta change. The symbols below each imprinted region highlight the mechanisms defined in (A). (C) Ideograms as in (B) for the ExE lineage. (D) Summary of X-linked regulator dependent and independent genes, as well as genes that escape the process of imprinted XCI. The model on the right illustrates the mechanism for how the inactive X is maintained in a silent state by *Xist* and Polycomb.

The extraembryonic lineage shares regulation for the majority of these ICR-regulated regions (*n* = 11) but includes additional ICR-controlled lncRNA clusters, as well as seven non-canonical G9a regulated imprinted regions (**Figure 5C**). Two lncRNA controlled clusters are regulated by *Kcnq1ot1* and *Airn* and incorporate known and novel secondary targets, all located in previously defined extraembryonic-specific silencing domains (Andergassen et al., 2017, 2019; Schertzer et al., 2019). For both clusters, we observe strong dependency for PRC and, to a lesser degree, for G9a, in line with previous reports where these regulators establish repressive chromatin at target loci (Nagano et al., 2013; Pandey et al., 2008; Terranova et al., 2008; Wagschal et al., 2008). The G9a-dominant imprinted regions include known and novel non-canonical imprinted genes and mostly lie in defined clusters, such as the *Smbt2, Jade1*, and *Slc38a4* imprinted regions and ERV-driven gene loci (**Figure 5C**). Finally, our data provide an improved understanding of imprinted XCI, including lncRNA Xist-based PRC2 recruitment that is most important for maintaining suppression at regions distal to primary *Xist* targets (**Figure 5D**).

## DISCUSSION

In this study, we systematically perturbed multiple repressive pathways in highly polymorphic embryos to investigate the scope of epigenetically maintained parental imprinting. We find that most canonically imprinted genes are shared between the embryonic and extraembryonic lineages. In contrast, non-canonical imprinting is largely restricted to the extraembryonic lineage. For instance, we identify a new H3K9 methylation-based mechanism that suppresses maternal ERV-driven promoters, which could also represent a more general strategy for gene regulation outside of imprinting. Previously, oocyte-specific H3K27me3 was described as a mechanism for non-canonical imprinting within preimplantation embryos (Inoue et al., 2017c). However, our systematic investigation across multiple different regulators finds only a slight effect in zygotic PRC2 mutants on the allelic ratios of these targets, but complete loss of imprinting and associated reactivation in *G9a* mutants, suggesting H3K9me is critical for establishing non-canonical imprinting.

We also provided mechanistic insides of long-range silencing by imprinted lncRNAs in extraembryonic lineages. Previous reports observed that *Kcnq1ot1* and *Airn* are expressed from the zygote stage and deployed to local targets to recruit PRCs (Andergassen et al., 2017, 2019; Nagano et al., 2008; Schertzer et al., 2019; Terranova et al., 2008). Notably, we find that *Dnmt1* KO embryos activate the silent *Kcnq1ot1* allele but do not affect imprinted expression of its PRC2-dependent targets. To capture both forms of deregulation independently, a transient window in preimplantation may allow lncRNAs to relay germline ICR information to their genetic neighborhood before DNA methylation is lost. As such, both DNA methylation-specific ICRs and their secondary, PRC2-dependent targets must be established very soon after fertilization. This model would also explain why Dnmt1-dependent imprints are shared between embryonic and extraembryonic lineages. Finally, *Kcnq1ot1* targets in PRC mutants show biallelic expression, hinting that oocyte-specific PRC is sufficient to initiate imprinted silencing, but zygotic PRC is required to maintain it. A similar mechanism may apply for the novel maternally expressed imprinted genes *Pnldc1* and *T* that lie within the lncRNA *Airn* silencing domain, although *Airn* itself is only lowly expressed during this developmental period (Andergassen et al., 2017, 2019; Schertzer et al., 2019).

Furthermore, our approach allowed us to shed light on the XCI maintenance *in vivo*. We elucidated both PRCs as critical factors that maintain portions of the inactive X chromosome in a repressed state, while perturbing of DNA methylation or H3K9me pathways had no impact. Finally, we identified PRC-independent gene clusters on the X chromosome, which resembles the early binding sites of Xist-mediated suppression. This would suggest that different territories on the X are uniquely dependent on direct binding via *Xist* or translate this local cue to distal areas by Polycomb. Together, our data provide a comprehensive inventory of the epigenetic mechanisms of parental-specific imprinting, which is also fundamental for many X-linked diseases as well as imprinting disorders where unlocking the silent healthy allele is an attractive therapeutic strategy.

Notably, our study provides a platform for future investigation into the molecular genetics of parental imprinting and X inactivation by combining genetic perturbation with fertilization using highly polymorphic strains. For example, we establish that non-canonical imprinting requires G9a to maintain maternal silencing of ERV promoter-containing genes but do not yet understand the nature of imprint establishment in the maternal germline. Conditional knockouts in the female germline will further expand our understanding of the relationship between the epigenetic machinery that encodes these imprints and those that interpret them to facilitate maintenance. Similarly, our PRC mutants exhibit limited reactivation of autosomal imprints or of paternal X reactivation, with only some leaky expression, whereas *Dnmt1* and *G9a* exhibited little or no effect. It is possible that these regulators provide additional levels of repression that can only be observed when PRCs are absent. Double or triple regulator mutant strategies would permit an exact, quantitative investigation for each independent regulator’s contribution to chromosome-level silencing.

Upon completion of this work, it appears that very few, if any, parent-specific allelic expression cannot be explained with one of the three reported mechanisms described above. However, little is known about how these non-canonical imprints are innovated over evolutionary time or the degree to which they are conserved across eutherian mammals in comparison to classic ICRs. Moreover, their striking enrichment in the placental lineage further highlights this tissue as a domain for expanded epigenetic innovation in mammals. Future studies in these areas will provide greater clarity for the roles these additional imprinting mechanisms play in supporting fetal development.

## FIGURES

**Figure S1.**
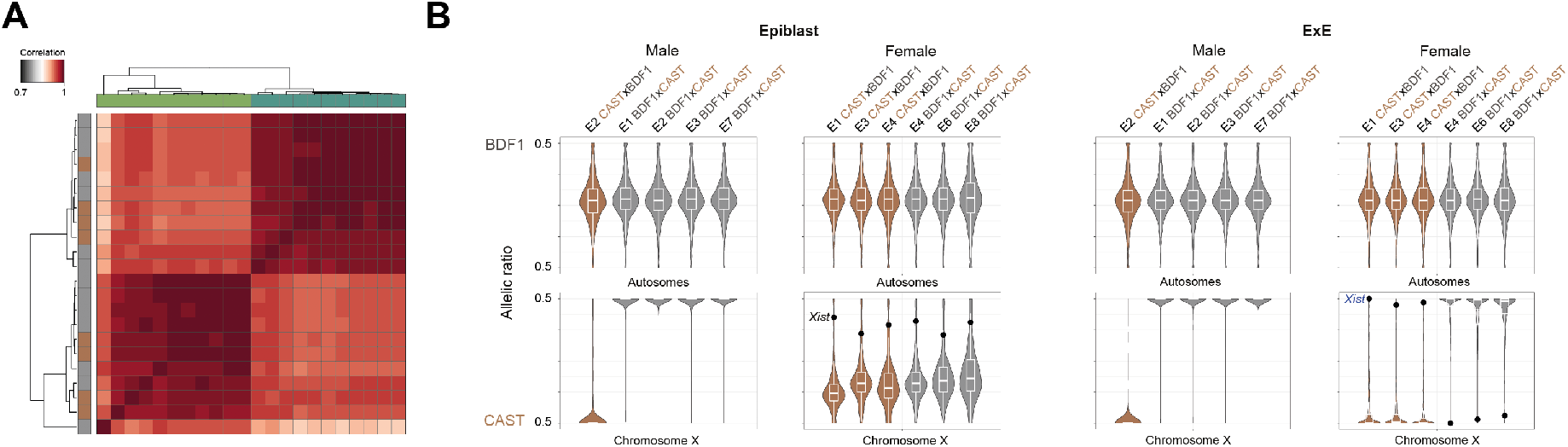
(A) Unsupervised clustering of pairwise Pearson correlation from log-transformed expression levels (TPM) of the forward (BDF1xCAST) and reverse (CASTxBDF1) cross confirms the lineage identity. (B) Violin plots displaying the allelic ratio of autosomal and X-linked genes from forward and reverse crosses for both genders in the Epiblast and ExE. The black dot highlights the lncRNA *Xist*.

**Figure S2.**
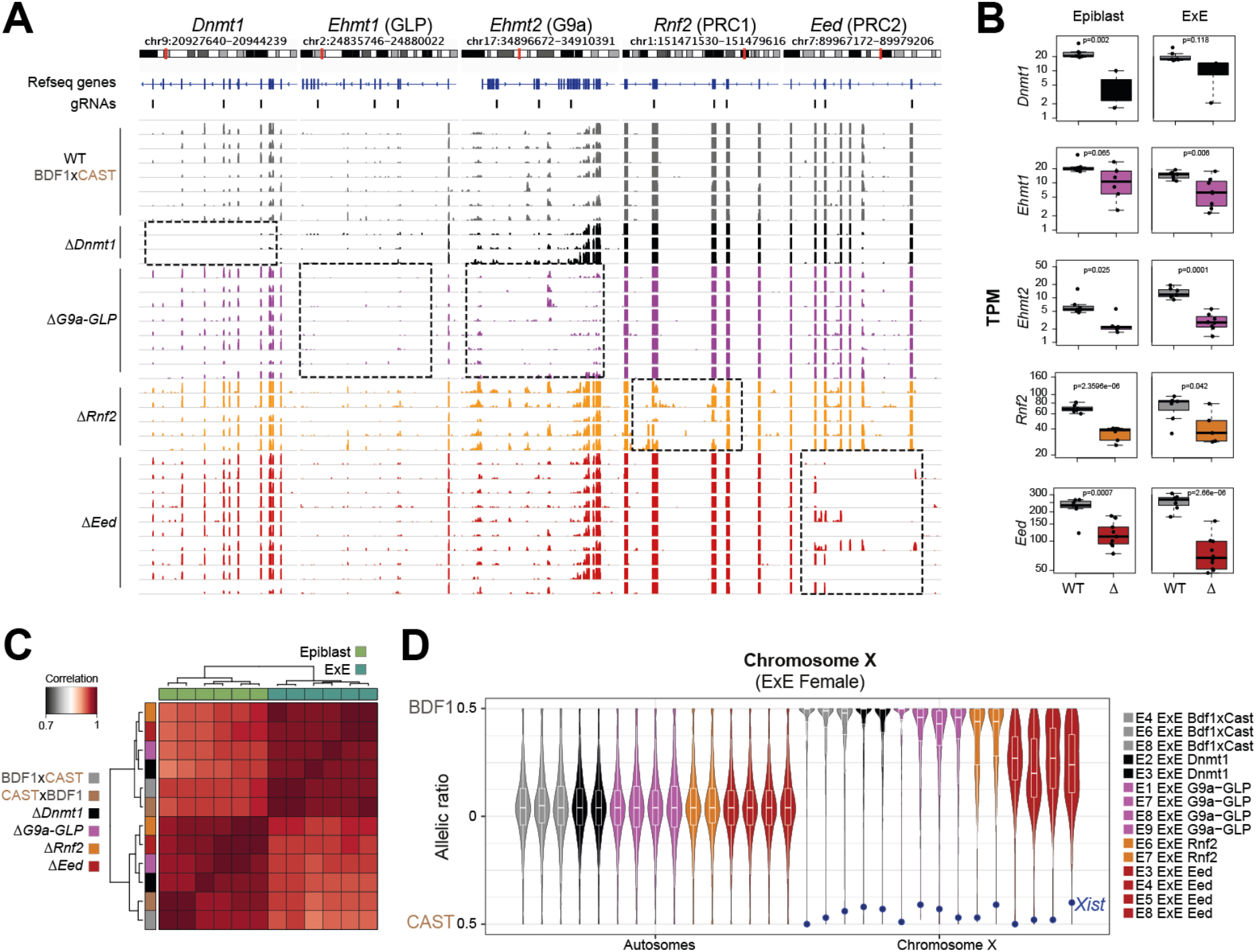
(A) Genome browser tracks showing the ExE RNA-seq results and gRNA positions of the targeted loci *Dnmt1, Ehmt1, Ehmt2, Rnf2*, and *Eed* for wildtype and regulator disrupted embryos. (B) Boxplot showing the expression levels (TPM) of the targeted loci between wildtype and KO embryos for the Epiblast and ExE lineage. (C) Unsupervised clustering of pairwise Pearson correlation from wildtype and regulator disrupted samples show broad retention of epiblast and ExE transcriptional signatures (TPM, replicates are combined using the average). (D) Violin plots displaying the allelic-ratio of X-linked genes from WT and regulator knockouts for every female ExE replicate (left). The blue dot highlights the allelic ratio of the lncRNA *Xist*.

**Figure S3.**
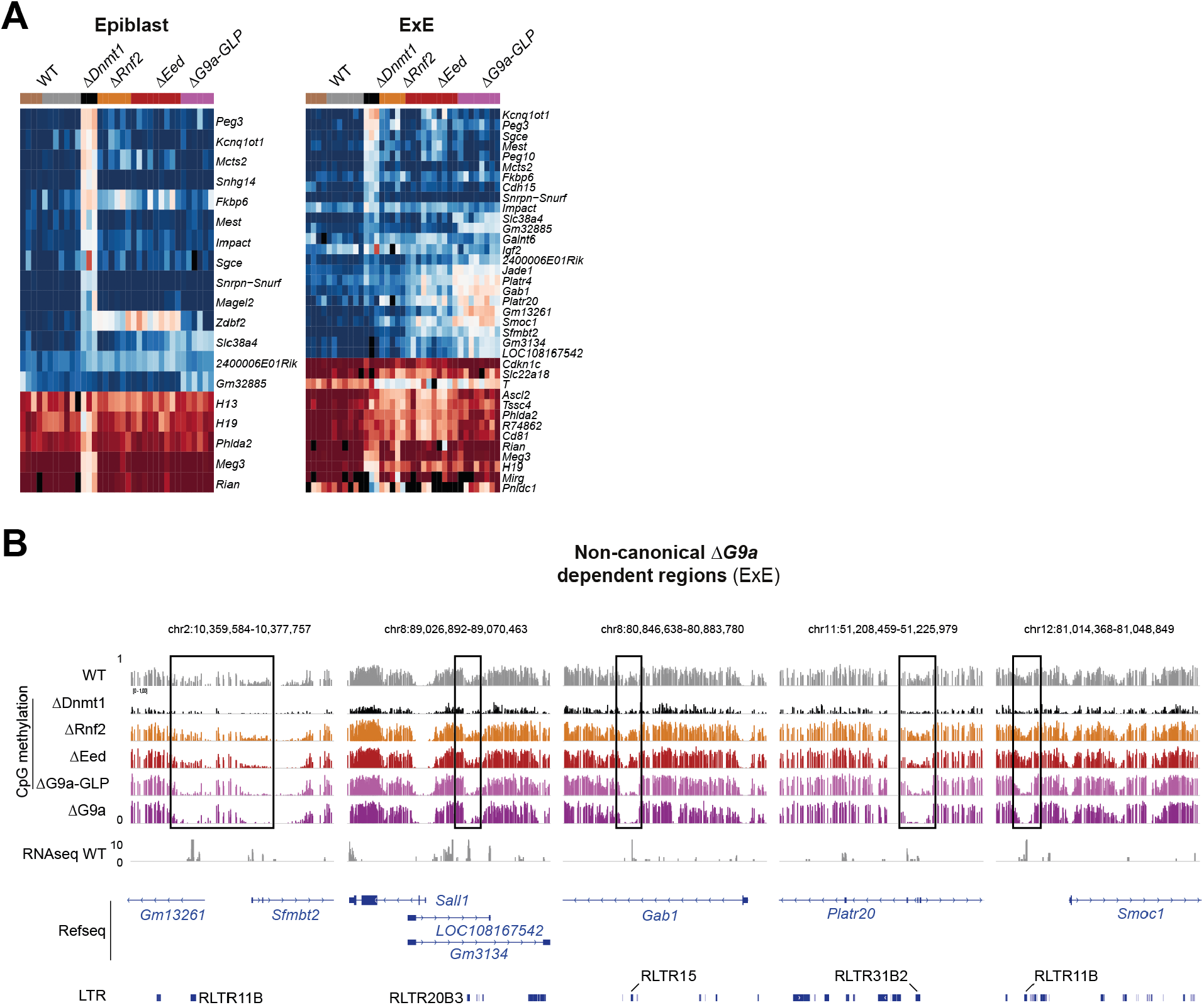
(A) Allelic ratio of imprinted genes between wildtype (BDF1xCAST) and regulator disruption identified in the epiblast (left) and ExE (right) lineage for each embryo. Heatmap ranked by delta allelic ratio change between WT and *Dnmt1* deletion. The black color indicates non-informative allelic ratios for a given gene and replicate. (B) Genome browser tracks for WGBS and RNA-seq data from WT and regulator disrupted E6.5 ExE lineages capturing the non-canonical G9a dependent imprinted loci *Gm13261, Sfmbt2, LOC108167542, Gm3134, Gab1, Platr20, and Smoc1*. Boxes highlight G9a dependent hypomethylated DMRs (Overlapping ERV LTRs are indicated).

**Figure S4.**
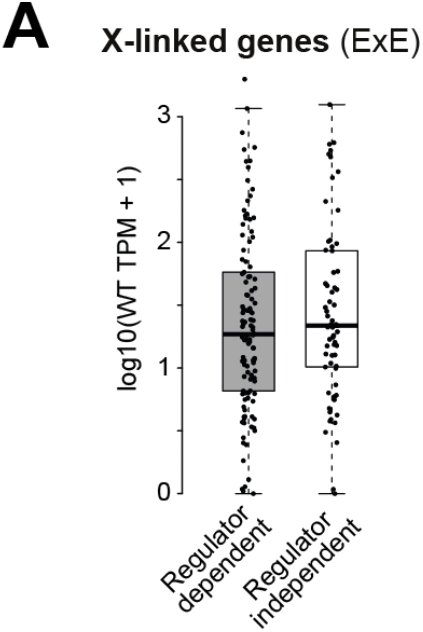
(A) Wildtype expression level (TPM) of regulator dependent and independent X-linked genes in ExE.

## METHOD DETAILS

### Mouse strains

All procedures have been performed in our specialized facility, followed all relevant animal welfare guidelines and regulations, and were approved by Harvard University IACUC protocol (28-21). CAST/EiJ (CAST) and B6D2F1/J (BDF1) mice were purchased from the Jackson Laboratory.

### Disrupting epigenetic regulators in F1 background using zygotic CRISPR–Cas9 injection

Embryos were generated as previously described (Wang et al., 2013). Briefly, BDF1 strain female mice (age 6–8 weeks, Jackson Labs) were superovulated by serial Pregnant Mare Serum Gonadotropin (5 IU per mouse, Prospec Protein Specialists) and human chorionic gonadotropin (5 IU, Millipore) injections 46 h apart. The following day, MII stage oocytes were isolated in M2 media supplemented with hyaluronidase (Millipore) and stored in 25 μl drops of pre-gassed KSOM with half-strength concentration of amino acids (Millipore) under mineral oil (Irvine Scientific). Zygotes were generated by piezo-actuated intracytoplasmic sperm injection (ICSI. See Ref. (Grosswendt et al., 2020)) using thawed CAST strain sperm in batches of 30–50 oocytes and standard micromanipulation equipment, including a Hamilton Thorne XY Infrared laser, Eppendorf Transferman NK2 and Patchman NP2 micromanipulators, and a Nikon Ti-U inverted microscope. Alternatively, for the reciprocal cross, BDF1 males were naturally mated to CAST females and screened for copulation plugs, after which E6.5 stage embryos were isolated accordingly, with the date of the copulation plug score day E0.5.

For zygotic disruption, pronuclear stage 3 (PN3) zygotes were injected with 200 ng μl^−1^ *Cas9* mRNA and a 100 ng μl^−1^ equimolar ratio of 3–4 sgRNAs targeting different exons of an epigenetic regulator gene locus (designed using ChopChop (Labun et al., 2016) and the IDT CRISPR–Cas9 guide RNA checker, as previously described in Ref. (Smith et al., 2017)). Injections utilized the same microinjection setup and Piezo-actuated injection of front-filled 6-7 μm injection needles. At around 84 h after fertilization, cavitated blastocysts were transferred into the uterine horns of pseudopregnant CD-1 strain females (25–35g, Charles River) generated by mating with vasectomized SW strain males (Taconic), which results in a 24 h offset in gestational time to accommodate implantation, after which animals were monitored for 5 days for embryo isolation at E6.5.

### Embryo isolation and Library preparation

At E6.5, animals were euthanized and the uterine horn removed. Purified epiblast and ExE were isolated according to Ref. (Chenoweth and Tesar, 2010) with a few modifications. Briefly, E6.5 embryos were removed from deciduae and transferred to independent 25 μL drops of M2 media. Using a glass flame-pulled capillary, Reichart’s membrane was reflected, and the embryo carefully bisected along the epiblast, ExE boundary. Then, each Epiblast/ExE pair was transferred into an individual drop of dissociation medium containing 0.5% trypsin and 2.5% pancreatin in PBS (w/v, Sigma). Embryos were cultured with slow orbital rotation at 4 ° C for 15 minutes, after which they were transferred into new M2 drops. After ∼5 minutes of resting, Epiblast and ExE were passed through a slightly narrower flame-pulled glass capillary to remove the visceral endoderm without disrupting the target tissue. Finally, each tissue was serially washed in 0.1% BSA in PBS prior to snap freezing in lysis buffer.

Approximately 300-600 cells were collected from E6.5 embryos and directly transferred to 2.6 µl of Lysis Buffer (Takara Bio USA, Inc.) followed by snap-freezing at -80°C in preparation for cDNA synthesis using the SMART-Seq v4 assay. Full-length cDNA was prepared using the SMART-Seq v4 Ultra Low Input RNA Kit for Sequencing and sequencing libraries prepared using the Nextera XT DNA library preparation kit (Illumina). The resulting libraries were evaluated using a 4200 TapeStation (Agilent Technologies) and quantified by qPCR. Libraries were pooled and sequenced on an Illumina NovaSeq SP or S1 flow cell using paired-end, 50 bp reads.

### RNA-seq and allele-specific analysis

The RNA-seq data were aligned to the mm10 reference genome using the STAR aligner (Dobin et al., 2013) (STAR version 2.5.0 c: –outFilterMultimapNmax 1). The read counts for every RefSeq isoform (RefSeq gene annotation (downloaded February 2018) were determined using the htseq-count phyton script (Anders et al., 2015) (version 0.6.1) and further normalized in R.

The Allelome.PRO approach was used to calculate allele-specific expression from the RNA-seq data as outlined in (Andergassen et al., 2015). Briefly, the Allelome.PRO pipeline uses strain-specific single-nucleotide polymorphisms (SNPs) to assign RNA-seq reads to the corresponding allele in F1 crosses. To obtain allele-specific ratios and scores from a single replicate, we slightly modified the pipeline (We will upload the modified Allelome.PRO version on SourceForge upon paper acceptance: https://sourceforge.net/projects/allelomepro/). The Allelome.PRO approach requires a gene and SNP annotation:

For the gene annotation, we used RefSeq, including 35856 genes. To avoid unreliable allele-specific calls, we modified the annotation for known overlapping imprinted genes: First, we removed all the isoforms of *H13, Kcnq1*, and *Copg2* overlapping the genes *Mcts2, Kcnq1ot1*, and *Mest2*, respectively. Second, we removed the two genes *Peg3os* and *Gm33149*, which overlap the known imprinted genes *Peg3* and *Gm32885*, respectively. Third, we truncated the gene *Tsix* to avoid overlap with *Xist* and *Gm32061* to prevent overlap with the two imprinted genes *Ndn* and *Magel2*. Finally, we assigned the gene name *Snrpn-Snurf* to the *Snrpn* and *Snurf* isoforms since both belong to the same gene.

To generate the SNP annotation for our F1 crosses between CAST and BDF1 (BDF1: F1 cross between C57BL/6J female x DBA/2J), we first derived 20,606,390 high confidence SNPs between CAST and C57BL/6J and 20,507,026 between CAST and DBA/2J (Keane et al., 2011). From the two SNP annotation files, we only used SNPs where the C57BL/6J allele was shared between DBA/2J (shared SNP nr: 17,967,587). Finally, we used only exonic SNPs, resulting in a final number of 1,513,184 SNPs. The Allelome.PRO “minread” parameter was set to 1 to include SNPs covered by one read. Genes with less than 10 read overlapping SNPs have been assigned to non-informative.

### Whole-genome bisulfite sequencing (WGBS) analysis

The E6.5 ExE *ΔG9a-Glp* sample was isolated and processed into WGBS libraries using the Accel-NGS Methyl-seq kit as described in (Grosswendt et al., 2020). We utilized our previously generated ExE WT (*n* = 2), *ΔG9a* (*n* = 1) samples (Grosswendt et al., 2020) together with the generated *ΔG9a-GLP* sample to define G9a specific differentially methylated regions (DMRs) in E6.5 ExE. CpGs with less than ten reads were removed for the downstream analysis. CpGs were binned over a 1kb window, filtering out all windows with less than 10 CpG. Window methylation levels were combined using the average. Next, we averaged WT and G9a knockout *(ΔG9a* and *Δand GLP*) methylation levels and calculated the delta (WT-KO). *Δ me* ExE DMRs were defined by havening a minimum difference of 0.2 to WT (|delta cutoff| ≥ 20%). Feature enrichment of the identified DMRs over background was calculated for intergenic DMRs, genic (±1kb of TSS, RefSeq annotation), and different repeat classes (RepeatMasker tracks, UCSC) using the Fisher’s exact test.

## ACKNOWLEDGEMENTS

RNA sequencing and library preparation were performed at the Bauer Core Facility at Harvard University. AM was supported by NIH grants (DP3K111898 and P01GM099117) and the Max Planck Society.

## DATA AND CODE AVAILABILITY

All datasets have been deposited in the Gene Expression Omnibus (GEO) and are accessible under GSE171206. Previously published data used in this study include WGBS data from GSE137337 for E6.5 ExE WT, *ΔDnmt1, ΔRnf2, ΔEed*, and *ΔG9a*.

